# SimpleP2G: Importance of Image Encoders for Gene Expression Prediction from Pathology Images

**DOI:** 10.64898/2026.11.23.768493

**Authors:** Jinghao Wang, Jingjie Zhang, Chang-Yu Hsieh, Chunbin Gu, Pheng-Ann Heng

## Abstract

Pathology-image-to-gene-expression (P2G) studies often change the prediction head, image encoder, and spatial context together, making performance differences difficult to attribute. We examine these factors through paired image-input controls and explicit input regimes. Our main comparison covers five HEST-1K datasets, two 200-gene panels, and three selected held-out slides per dataset, using validation-selected checkpoints. Across all five datasets, replacing conventional image inputs with Virchow2 improves mean PCC-200 in 39 of 40 method–dataset–panel comparisons, while one comparison decreases. We introduce *SimpleP2G*, a direct gene-query decoder over frozen embeddings, and *SimpleP2G-cross*, which adds spatially positioned neighboring embeddings. With shared Virchow2 features, SimpleP2G has the highest single-patch mean PCC-200 on all five datasets under both panels; the cross-patch variant leads on four of five datasets under each panel. Decoder and context ablations show that greater depth and broader context do not consistently improve prediction. Together, these results suggest improving image representations before adding prediction-head complexity: a direct decoder over frozen pathology embeddings already provides a strong baseline. We hope our reimplemented image-encoder-upgraded baselines and SimpleP2G models serve as transparent, reproducible, and strong controls for future P2G research. The implementation is hosted at https://github.com/king159/SimpleP2G.

## 1 Introduction

H&E histology is routinely collected in tissue studies and clinical pathology, whereas spatial transcriptomics (ST) provides costly molecular measurements at aligned tissue locations. Predicting spatially resolved gene expression from H&E images—pathology-image-to-gene-expression (P2G) prediction—therefore offers a practical way to connect abundant morphology with lower-throughput molecular profiling.

Recent P2G systems have invested heavily in the prediction stage: contrastive retrieval, diffusion or flow generation, gene-ranking objectives, and spatial graph or Transformer architectures. Yet these methods frequently differ in more than the proposed head. A newer system may also receive features from a stronger pathology foundation model, use neighboring patches or whole-slide context, and follow a different training schedule. Attributing such a compound improvement to head design alone can be misleading. We therefore ask how much can be gained by improving the visual representation while retaining the downstream prediction method.

Our paired comparisons across all five datasets show positive mean PCC-200 changes in 39 of 40 method–dataset–panel combinations. For example, replacing HisToGene’s raw-patch input with Virchow2 features increases its three-slide mean HER2ST HMHVG PCC-200 from 0.1209 to 0.4321. BLEEP on Brain HMHVG instead decreases from 0.3376 to 0.3258, motivating within-method comparisons that hold the split, objective, and training schedule fixed. To make the comparison concrete, we introduce **SimpleP2G**, a direct baseline. A frozen image encoder produces one patch embedding; a small MLP converts it to conditioning tokens; and learnable gene queries cross-attend to those tokens before predicting expression. There is no trainable image backbone, retrieval bank, or iterative sampling procedure. For cross-patch prediction, **SimpleP2G-cross** keeps the decoder and loss unchanged while expanding the conditioning sequence to include spatially positioned neighboring patches and, optionally, a slide-summary token. This design provides a transparent reference for assessing how much complexity is needed after choosing a strong image representation.

Our main experiments compare ten models on Brain, PRAD, HER2ST, Lung, and ccRCC under two gene panels, with separate single-patch and cross-patch settings. Every method uses frozen Virchow2 image features and the same three selected test-slide folds, training-only gene panels, and validation assignments. Both settings therefore share the same dataset coverage. Decoder and context ablations use one fold per dataset to examine depth, objective, neighborhood size, and slide information separately.

We summarize our contributions as follows:

- We quantify the effect of replacing image representations within existing prediction heads. All 120 paired folds are evaluated, giving 39 positive and one negative mean change across 40 method–dataset–panel combinations.
- We introduce SimpleP2G and SimpleP2G-cross, direct genequery decoders over frozen image embeddings. They provide competitive references in both input settings, while ablations identify cases where simpler heads or less context perform better.
- We compare eight baseline implementations and our two models across five HEST-1K datasets and two gene panels, reporting variation across three held-out slides. Gene-DML retains its target, neighbor, and global pathways in the cross-patch setting; STRank and M2TGLGO adaptations are documented explicitly.

## 2 Related Work

### 2.1 Target-Patch and Cross-Patch P2G

Early work established that local morphology contains predictive signal for spatial expression. ST-Net [7] treats each measured location as an image-to-gene regression example, whereas HisToGene [19] lets spots from the same tissue section interact through a spatially informed Transformer. These works define two distinct information settings: a single-patch predictor observes only the patch centered at the queried spot; a cross-patch predictor also observes other image regions from the section. We retain this distinction throughout because an architectural comparison is meaningful only when the available image evidence is matched.

Later single-patch methods replace direct regression with more elaborate objectives. BLEEP [31] learns a shared image–expression space through contrastive training and predicts expression by retrieval. Stem [35] uses conditional diffusion, GenAR [18] uses autoregressive generation, and STRank [16] emphasizes relative gene ordering. These objectives impose useful inductive biases, but they may also restrict how encoder information reaches the prediction: retrieval is bounded by a reference bank, while iterative generators add a costly inference path. Our single-patch experiments ask whether their reported advantage remains after their image representation is upgraded.

Recent research increasingly exploits larger spatial context. Gene-DML [23] combines target, neighbor, and global image pathways with multi-level discrimination and therefore belongs to the cross-patch setting. STFlow [9] performs whole-slide flow matching with spatial attention. MERGE [6] constructs a hierarchical graph from spatial and feature-space relations, and M2TGLGO [21] combines spatial topology with gene-level relations. HistoPrism [8] and FmH2ST [30] further explore slide- or foundation-model-level information. Such methods may legitimately outperform target-only predictors because they solve a richer problem. We therefore compare them in the cross-patch setting and, where their inter-faces permit, evaluate both their released/default image input and Virchow2. SimpleP2G-cross uses an explicit target-plus-neighbor conditioning sequence; baselines that natively consume the full section retain that larger context, so the simple control is not given an information advantage.

### 2.2 Image Representation versus Prediction Head

Pretrained representations are increasingly used for medical image understanding [3, 4] and molecular prediction [33, 34]. In pathology,UNI [2], CONCH [13], Virchow/Virchow2 [29, 36], and Hibou [15] learn from large histology collections; DINOv2 [17], DINOv3 [22], and SAM [11] provide strong general-vision controls.

In P2G, image representations are often treated as interchangeable implementation details, even though changing them can alter both prediction accuracy and the behavior of a downstream method. The issue is especially important for feature-dependent heads: nearest-neighbor retrieval changes when the embedding geometry changes, and a graph method may construct different edges or clusters from a new feature space. Accordingly, we distinguish two questions. An *encoder upgrade* keeps a method’s downstream computation fixed and measures how much it benefits from a stronger input representation. A *head comparison* fixes the encoder and accounts explicitly for the context available to each method; when a native whole-slide method uses more evidence, we retain that advantage rather than claiming an exactly matched input. This two-step comparison supports a more defensible claim than either an unpaired leaderboard or the assertion that encoder quality always dominates.

## 3 Method

### 3.1 Problem Formulation

A tissue section contains aligned ST spots S = {*s*_1_, …, *s*_*M*_}. Spot *s*_*i*_ has an H&E patch v_*i*_ ∈ ℝ^*L*×*L*×3^, spatial coordinate r_*i*_ ∈ ℝ ^2^, and expression vector x_*i*_∈ ℝ^*C*^ for a selected panel of *C* genes. Our models train on log_2_ (count + 1) expression and predict 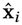 without observing expression at test time.

We evaluate two information settings. A single-patch method predicts

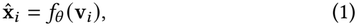

whereas a cross-patch method may also use image-only context

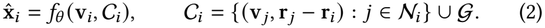

Here *N*_*i*_ is a spatial neighborhood and *G* denotes optional sectionlevel image information. Gene expression from neighbors is never an input. Since the second setting contains strictly more image evidence, claims about head quality are made only within a fixed setting and, for paired comparisons, with a fixed encoder.

### 3.2 SimpleP2G

Let *ϕ* be a frozen image encoder and 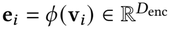 its singlepatch embedding. In the main experiments *ϕ* is Virchow2 [36], represented by its class token concatenated with the mean patch token (*D*_enc_ = 2560). The trainable predictor is deliberately restricted to expression decoding:

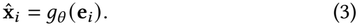

#### Image conditioning

A two-layer MLP with SiLU activation maps e_*i*_ to*T* conditioning tokens 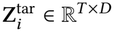, with*T* = 4 by default. This keeps the decoder interface fixed across encoder dimensions.

#### Cross-patch conditioning

For SimpleP2G-cross, *N*_*i*_ contains the *K* = 24 nearest measured spots excluding *i*. The same frozen encoder *ϕ* produces target and neighbor embeddings. Each neighbor is projected to one token and receives a learned encoding of its relative coordinate, normalized by the section’s median nearest-neighbor spacing:

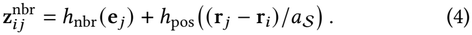

Missing neighbors are padded and masked. An optional slide token is 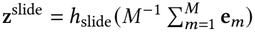. The decoder attends to the concatenation of whichever sources are enabled:

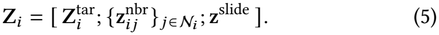

Thus target only, neighbors only, target plus neighbors, and target plus neighbors plus slide differ only in the conditioning sequence.

#### Gene-query decoder

A learnable table Q ∈ ℝ^*C*×*D*^ contains one query per predicted gene. Queries pass through *L* pre-norm Transformer decoder layers with self-attention among genes, cross-attention to Z_*i*_, and a gated feed-forward block with residual connections and RMSNorm [32]. Unless stated otherwise, *L* = 12, *D* = 384, with six attention heads; dropout follows Appendix A. Each final query predicts a mean 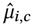 and log-variance log 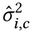; softplus makes the mean non-negative in transformed expression space.

#### Loss and Optimization

We train both variants with the same Gaussian negative log-likelihood:

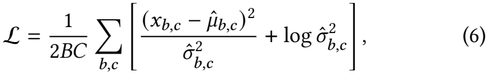

where log 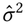 is clamped to [−10, 10]. We use AdamW with *β* = 0.9, 0.999, gradient clipping at 1.0, linear warmup, and cosine decay where specified. Both variants require one deterministic forward pass at inference.

## 4 Experiments

### 4.1 Datasets

Our main evaluation uses five public spatial transcriptomics datasets from HEST-1k [10]: Brain, PRAD, HER2ST, Lung, and ccRCC. They span five organs, human and mouse tissue, and three sequencing platforms (Visium, Spatial Transcriptomics v1, and Xenium), comprising 117 tissue sections and 226,656 measured spots. We evaluate three selected leave-one-slide-out folds per dataset. Each fold reserves one additional slide for validation and trains on the remaining slides. The assignments are fixed before training and shared by all methods; these are slide-level rather than patient-level holdouts.

### 4.2 Experimental Setup

#### Gene Selection

Following previous works [19, 35], we construct two 200-gene panels after filtering mitochondrial and ribosomal genes: *HMHVG-200*, genes with high mean and high variance, and *HVG-200*, batch-aware highly variable genes selected across slides.

Panels are selected independently from each fold’s training expression, excluding validation and test slides. Evaluation targets are log_2_(count+1).

#### Evaluation Metrics

We report Pearson correlation coefficient (PCC), MSE, and MAE. PCC-200 averages per-gene correlations over the selected panel and is the primary metric. PCC-50 averages the highest 50 per-gene correlations as a supplementary summary. Genes with constant observed expression are excluded identically for all methods; constant predictions for variable targets receive zero correlation. Every method uses the same held-out spots and gene panel. Main results report means and sample SDs across the three slides with seed 42; ablations use fold 0 and the same seed. Metrics are rounded to four decimals. Per-fold main results and paired encoder scores are provided as CSV files in the code repository.

#### Baselines and Input Regimes

Single-patch methods receive only the queried patch; cross-patch methods may also use surrounding information. The single-patch comparison includes BLEEP [31], Stem [35], and STRank^†^ [16]. The cross-patch comparison includes Gene-DML [23], HisToGene [19], STFlow [9], MERGE [6], and M2TGLGO [21]. Reimplementation and adaptation details are given in Appendix B.

### 4.3 Implementation Details

#### Models

Main comparisons and decoder/context ablations use a shared frozen Virchow2 encoder for every available image input. SimpleP2G uses its 2560-dimensional embeddings, a 12-layer decoder of width 384, six attention heads, and four target-conditioning tokens. SimpleP2G-cross adds 24 spatial nearest neighbors; relative coordinates are normalized by median spot spacing. Unless specified otherwise, SimpleP2G-cross uses target and neighbor inputs without a slide token. The optional slide token averages same-slide spot embeddings.

#### Training

All methods train for 100 epochs except Stem, which uses 200. A separate validation slide selects the checkpoint with the highest recorded validation PCC-200. Baseline learning rates follow their released implementations; SimpleP2G’s dataset-specific settings are listed in Appendix A. SimpleP2G-cross uses learning rate 5 × 10^−5^, weight decay 0.05, and dropout 0.05 throughout. Both SimpleP2G variants use AdamW [12], gradient clipping at 1.0, and EMA. Cross-patch training samples slide-level chunks of up to 256 targets, while single-patch training shuffles pooled-spot batches.

#### Batching and Checkpoints

In SimpleP2G, pooled batches contain 256 spots and use an augmentation ratio of 2. For SimpleP2G-cross, each slide contributes eight chunks per epoch, with the target, neighbors, and optional slide summary drawn from that slide. A slide is the batch item for slide-based baselines. M2TGLGO caps this item at 1,024 spots, while BLEEP retrieves training examples from its expression bank. These batching choices preserve the native units required by the different heads. The training split supplies expression targets and any training bank. Validation determines only the checkpoint. Ties select the earliest epoch, and each selected checkpoint is evaluated once on the corresponding test slide without refitting.

### 4.4 Strong Impact of Image Encoder

Table 1 replaces the image representation in four prediction heads with Virchow2. Reference inputs are ImageNet-pretrained ResNet50 for BLEEP, raw patches for HisToGene, features from MERGE’s released TCGA-pretrained ResNet18 checkpoint, and ImageNet-pretrained ResNet50 following M2TGLGO’s released backbone choice. Each pair retains the downstream head, objective, split, and training schedule while adapting the input projection. Retrieval neighbors and feature-derived graph structure are recomputed when required. Arrows report three-slide means with SDs; Δ is the absolute paired mean PCC change.

**Table 1:** Impact of upgrading image encoders. Reference input → Virchow2 PCC-200 (mean ± sample SD over three held-out slides); Δ is the absolute paired mean change. The higher mean within each pair is bold. ^†^ denotes our adaptation; see Appendix B.

| **Method** | **HMHVG** |  | **HVG** |  |
|---|---|---|---|---|
|  | Reference → Virchow2 | ΔPCC | Reference → Virchow2 | ΔPCC |
| *Brain* [28] |  |  |  |  |
| BLEEP | **0.3376**±0.1311 → 0.3258±0.1773 | -0.0118 | 0.1408±0.0635 → **0.1488**±0.0559 | +0.0079 |
| HisToGene | 0.0338±0.0278 → **0.3240**±0.0759 | +0.2902 | 0.0244±0.0215 → **0.1157**±0.0071 | +0.0913 |
| MERGE | 0.1856±0.0501 → **0.3466**±0.1364 | +0.1610 | 0.0166±0.0713 → **0.1225**±0.0418 | +0.1059 |
| M2TGLGO<sup>†</sup> | 0.3381±0.2180 → **0.4585**±0.1292 | +0.1205 | 0.1526±0.0223 → **0.1831**±0.0506 | +0.0305 |
| *PRAD* [5] |  |  |  |  |
| BLEEP | 0.4218±0.1286 → **0.4791**±0.1209 | +0.0573 | 0.1458±0.0285 → **0.1668**±0.0333 | +0.0211 |
| HisToGene | 0.2942±0.0734 → **0.4839**±0.0859 | +0.1898 | 0.0796±0.0064 → **0.1664**±0.0187 | +0.0868 |
| MERGE | 0.3743±0.1280 → **0.4295**±0.1425 | +0.0552 | 0.0983±0.0247 → **0.1524**±0.0297 | +0.0541 |
| M2TGLGO<sup>†</sup> | 0.4467±0.0539 → **0.5596**±0.0928 | +0.1129 | 0.1596±0.0218 → **0.2078**±0.0216 | +0.0482 |
| *HER2ST* [1] |  |  |  |  |
| BLEEP | 0.4711±0.1900 → **0.4980**±0.1926 | +0.0269 | 0.2050±0.1304 → **0.2129**±0.1155 | +0.0079 |
| HisToGene | 0.1209±0.2053 → **0.4321**±0.2574 | +0.3113 | 0.1371±0.1088 → **0.1988**±0.1307 | +0.0617 |
| MERGE | 0.3940±0.2332 → **0.4637**±0.2160 | +0.0697 | 0.1533±0.1551 → **0.2093**±0.1368 | +0.0561 |
| M2TGLGO<sup>†</sup> | 0.4804±0.1910 → **0.5235**±0.1970 | +0.0431 | 0.2041±0.1195 → **0.2403**±0.1302 | +0.0362 |
| *Lung* [27] |  |  |  |  |
| BLEEP | 0.8123±0.0193 → **0.8401**±0.0139 | +0.0278 | 0.4239±0.0314 → **0.4268**±0.0242 | +0.0029 |
| HisToGene | 0.7651±0.0292 → **0.8171**±0.0212 | +0.0520 | 0.3686±0.0333 → **0.4124**±0.0303 | +0.0439 |
| MERGE | 0.8042±0.0055 → **0.8404**±0.0115 | +0.0362 | 0.3930±0.0124 → **0.4296**±0.0273 | +0.0366 |
| M2TGLGO<sup>†</sup> | 0.8419±0.0197 → **0.8708**±0.0105 | +0.0289 | 0.4348±0.0330 → **0.4675**±0.0207 | +0.0327 |
| *ccRCC* [14] |  |  |  |  |
| BLEEP | 0.1958±0.1468 → **0.3039**±0.1536 | +0.1081 | 0.0768±0.0329 → **0.0955**±0.0410 | +0.0187 |
| HisToGene | 0.1100±0.1083 → **0.3939**±0.0380 | +0.2839 | 0.0256±0.0269 → **0.1810**±0.0699 | +0.1554 |
| MERGE | 0.1372±0.0340 → **0.4205**±0.1367 | +0.2833 | -0.0183±0.0547 → **0.1475**±0.0652 | +0.1658 |
| M2TGLGO<sup>†</sup> | 0.3373±0.0606 → **0.4303**±0.1135 | +0.0930 | 0.1119±0.0270 → **0.1954**±0.0752 | +0.0835 |

Across 120 paired folds, Virchow2 improves the three-slide mean in 39 of 40 dataset–method–panel comparisons. Absolute mean changes range from −0.0118 to +0.3113. HisToGene on HER2ST HMHVG increases from 0.1209 to 0.4321, while BLEEP on HER2ST HVG increases from 0.2050 to 0.2129. The exception is BLEEP on Brain HMHVG, which decreases from 0.3376 to 0.3258. These comparisons demonstrate broad, method- and tissue-dependent representation gains.

The underlying slide pairs show how often a positive average agrees with the individual results. PCC-200 increases on 115 of 120 matched slides, with increases on all three slides in 35 of the 40 dataset–method–panel combinations. The five remaining combinations improve on two slides each. For example, the three changes for BLEEP on Brain HMHVG are +0.0645, +0.0241, and −0.1242. The negative mean for this combination reflects the magnitude of the third change. Across the paired runs, MSE decreases in 98 of 120 comparisons and MAE decreases in 96 of 120. These error metrics reflect expression magnitude, while PCC measures pattern agreement.

### 4.5 Target-Patch Results

Among the four single-patch methods in Table 2, SimpleP2G obtains the highest mean PCC-200 on all five datasets under both panels. Its five-dataset means are 0.5336 for HMHVG and 0.2485 for HVG; on Brain HMHVG it reaches 0.4365 _±0.1248_. Error-metric rankings remain mixed across methods. A direct decoder therefore provides a strong reference for specialized objectives and inference procedures under shared image features.

**Table 2:**
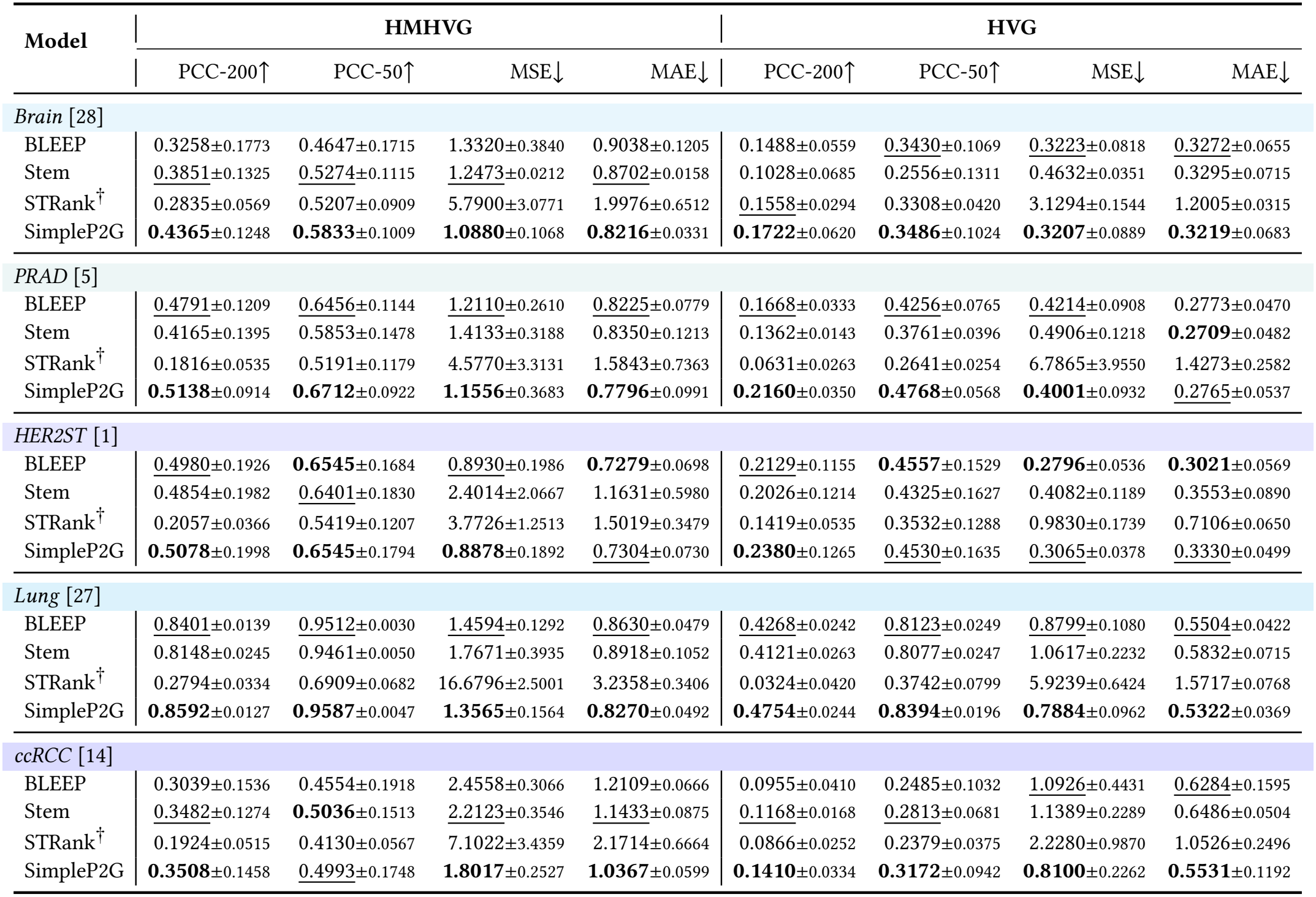
Single-patch results on five HEST-1K datasets. Mean±sample SD across three held-out slides, seed 42, with shared Virchow2 features and validation-selected checkpoints. Best and second-best means within this setting are bold and <u>underlined</u>. ^†^ denotes our adaptation; see Appendix B.

### 4.6 Cross-Patch Results

SimpleP2G-cross obtains the highest mean PCC-200 on four of five datasets under each panel in Table 3. M2TGLGO is strongest on PRAD HMHVG and ccRCC HVG. The five-dataset HMHVG/HVG means are 0.5789/0.2722 for SimpleP2G-cross, 0.5685/0.2588 for M2TGLGO, and 0.5538/0.2596 for Gene-DML. These results support competitive performance with tissue-specific exceptions; three selected slides provide descriptive variation rather than evidence of statistical superiority.

**Table 3:** Cross-patch results on five HEST-1K datasets. Mean ± sample SD across three held-out slides, seed 42, with shared Virchow2 features and validation-selected checkpoints. Best and second-best means within this setting are bold and <u>underlined</u>. ^†^ denotes our adaptation; see Appendix B.

| Model | HMHVG |  |  |  | HVG |  |  |  |
|---|---|---|---|---|---|---|---|---|
|  | PCC-200↑ | PCC-50↑ | MSE↓ | MAE↓ | PCC-200↑ | PCC-50↑ | MSE↓ | MAE↓ |
| *Brain* [28] |  |  |  |  |  |  |  |  |
| HisToGene | 0.3240±0.0759 | 0.4460±0.0674 | 1.4185±0.3742 | 0.9559±0.1416 | 0.1157±0.0071 | 0.3318±0.0205 | 0.3326±0.1188 | 0.3391±0.0647 |
| STFlow | 0.3288±0.1099 | 0.4555±0.1036 | 1.3507±0.2034 | 0.9218±0.0767 | 0.1318±0.0645 | 0.3019±0.1078 | 0.3636±0.1175 | 0.3326±0.0592 |
| MERGE | 0.3466±0.1364 | 0.4767±0.1267 | 1.3722±0.2210 | 0.9343±0.0668 | 0.1225±0.0418 | 0.3251±0.0999 | 0.3319±0.0848 | 0.3539±0.0630 |
| M2TGLGO<sup>†</sup> | 0.4585±0.1292 | 0.5947±0.1030 | 1.2493±0.2949 | 0.8642±0.1139 | 0.1831±0.0506 | **0.4023**±0.0855 | 0.3215±0.0755 | 0.3423±0.0521 |
| Gene-DML | 0.4612±0.1103 | 0.6015±0.1133 | **1.1557**±0.1759 | **0.8486**±0.0582 | 0.1765±0.0637 | 0.3545±0.1175 | 0.3322±0.0777 | 0.3258±0.0746 |
| SimpleP2G-cross | **0.4702**±0.1232 | **0.6093**±0.1112 | 1.2012±0.3377 | 0.8607±0.1158 | **0.1964**±0.0463 | 0.3934±0.0770 | **0.3120**±0.0848 | **0.3225**±0.0656 |
| *PRAD* [5] |  |  |  |  |  |  |  |  |
| HisToGene | 0.4839±0.0859 | 0.6238±0.0796 | 1.2872±0.3028 | 0.8764±0.1051 | 0.1664±0.0187 | 0.4460±0.0224 | 0.4011±0.0993 | 0.2961±0.0444 |
| STFlow | 0.4672±0.0848 | 0.6200±0.0767 | 1.3092±0.4776 | 0.8529±0.1583 | 0.2085±0.0239 | 0.4938±0.0210 | 0.3567±0.1179 | 0.2638±0.0562 |
| MERGE | 0.4295±0.1425 | 0.5996±0.1275 | 1.4461±0.5504 | 0.9274±0.1809 | 0.1524±0.0297 | 0.4265±0.0514 | 0.4050±0.1588 | 0.2978±0.0562 |
| M2TGLGO<sup>†</sup> | **0.5596**±0.0928 | **0.7092**±0.0871 | 1.2473±0.3640 | 0.8417±0.1068 | 0.2078±0.0216 | 0.5240±0.0357 | 0.3578±0.1020 | 0.2839±0.0506 |
| Gene-DML | 0.5076±0.1237 | 0.6777±0.1125 | 1.1477±0.3570 | 0.7841±0.0923 | 0.2275±0.0204 | 0.5264±0.0240 | 0.3570±0.1020 | 0.2695±0.0430 |
| SimpleP2G-cross | 0.5541±0.0974 | 0.7036±0.0899 | **1.0086**±0.3686 | **0.7312**±0.1162 | **0.2380**±0.0348 | **0.5288**±0.0414 | **0.3381**±0.1047 | **0.2532**±0.0511 |
| *HER2ST* [1] |  |  |  |  |  |  |  |  |
| HisToGene | 0.4321±0.2574 | 0.5744±0.2549 | 1.0042±0.2500 | 0.7850±0.0883 | 0.1988±0.1307 | 0.4216±0.1889 | 0.2892±0.0421 | 0.3276±0.0483 |
| STFlow | 0.4687±0.2277 | 0.6224±0.2107 | 0.9393±0.2166 | 0.7450±0.0781 | 0.2384±0.1430 | 0.4644±0.1795 | 0.2686±0.0470 | **0.2995**±0.0574 |
| MERGE | 0.4637±0.2160 | 0.6103±0.2036 | 0.9267±0.1318 | 0.7577±0.0396 | 0.2093±0.1368 | 0.4236±0.1902 | 0.2703±0.0442 | 0.3080±0.0506 |
| M2TGLGO<sup>†</sup> | 0.5235±0.1970 | 0.6671±0.1738 | 1.1374±0.5471 | 0.8191±0.2070 | 0.2403±0.1302 | **0.4839**±0.1727 | 0.2958±0.0776 | 0.3304±0.0832 |
| Gene-DML | 0.5159±0.1945 | 0.6661±0.1641 | **0.9084**±0.2115 | 0.7272±0.0748 | 0.2386±0.1323 | 0.4638±0.1793 | **0.2678**±0.0450 | 0.3010±0.0592 |
| SimpleP2G-cross | **0.5423**±0.1906 | **0.6892**±0.1592 | 0.9125±0.2829 | **0.7264**±0.1080 | **0.2460**±0.1221 | 0.4837±0.1648 | 0.2897±0.0329 | 0.3071±0.0553 |
| *Lung* [27] |  |  |  |  |  |  |  |  |
| HisToGene | 0.8171±0.0212 | 0.9492±0.0045 | 1.6274±0.1612 | 0.9222±0.0549 | 0.4124±0.0303 | 0.8039±0.0255 | 0.9087±0.0912 | 0.5790±0.0199 |
| STFlow | 0.8645±0.0113 | 0.9579±0.0052 | 1.3935±0.2484 | 0.8523±0.0755 | 0.4755±0.0230 | 0.8520±0.0185 | 0.7602±0.1010 | 0.5233±0.0294 |
| MERGE | 0.8404±0.0115 | 0.9448±0.0096 | 1.5162±0.0818 | 0.8918±0.0317 | 0.4296±0.0273 | 0.8242±0.0265 | 0.8272±0.0728 | 0.5669±0.0173 |
| M2TGLGO<sup>†</sup> | 0.8708±0.0105 | 0.9630±0.0057 | 1.4893±0.2314 | 0.9082±0.0903 | 0.4675±0.0207 | 0.8627±0.0191 | 0.8188±0.1409 | 0.5617±0.0512 |
| Gene-DML | 0.8754±0.0104 | 0.9638±0.0038 | 1.2407±0.2161 | 0.7885±0.0721 | 0.4855±0.0228 | **0.8699**±0.0192 | **0.7053**±0.1145 | 0.5151±0.0420 |
| SimpleP2G-cross | **0.8791**±0.0100 | **0.9667**±0.0058 | **1.1163**±0.1316 | **0.7395**±0.0482 | **0.4881**±0.0190 | 0.8655±0.0186 | 0.7210±0.0720 | **0.5006**±0.0259 |
| *ccRCC* [14] |  |  |  |  |  |  |  |  |
| HisToGene | 0.3939±0.0380 | 0.5356±0.0715 | 2.1641±0.7765 | 1.1353±0.1943 | 0.1810±0.0699 | 0.4007±0.1462 | 0.9471±0.5439 | 0.6209±0.1728 |
| STFlow | 0.4082±0.0843 | 0.5498±0.0958 | 1.8050±0.2944 | 1.0595±0.0981 | 0.1695±0.0632 | 0.3788±0.1330 | **0.7422**±0.2274 | 0.5588±0.1099 |
| MERGE | 0.4205±0.1367 | 0.5775±0.1481 | 1.7969±0.4232 | 1.0479±0.1053 | 0.1475±0.0652 | 0.3576±0.1446 | 0.9445±0.4028 | 0.6068±0.1648 |
| M2TGLGO<sup>†</sup> | 0.4303±0.1135 | 0.5848±0.1289 | 2.7646±0.9674 | 1.2979±0.2558 | **0.1954**±0.0752 | **0.4204**±0.1402 | 1.0222±0.3994 | 0.6499±0.1021 |
| Gene-DML | 0.4090±0.1610 | 0.5670±0.1845 | 1.7385±0.1159 | 1.0265±0.0199 | 0.1697±0.0570 | 0.3818±0.1360 | 0.7619±0.3262 | **0.5351**±0.1665 |
| SimpleP2G-cross | **0.4490**±0.1023 | **0.5979**±0.1227 | **1.7334**±0.3457 | **1.0167**±0.0888 | 0.1922±0.0655 | 0.4100±0.1363 | 0.8642±0.3564 | 0.5679±0.1355 |

### 4.7 Image Encoder Ablation

Table 4 compares six image encoders within SimpleP2G across all five datasets and both panels, using fold 0 and seed 42. Virchow2 has the highest PCC-200 in seven of ten combinations, including both panels on PRAD, HER2ST, and Lung. UNI leads on Brain HMHVG (0.3859 versus 0.3803 for Virchow2) and ccRCC HMHVG (0.3381 versus 0.2704). DINOv2-L leads on Brain HVG (0.1712 versus 0.1690). Thus, the encoder ranking depends on the tissue and panel, and a general image encoder can outperform pathology-specific encoders in an individual comparison.

**Table 4:** Image encoder ablation for SimpleP2G. PCC-200 on fold 0 of five datasets, both gene panels, seed 42, with validation-selected checkpoints. Best value per dataset/panel is bold.

| **Encoder** | **Brain** |  | **PRAD** |  | **HER2ST** |  | **Lung** |  | **ccRCC** |  |
|---|---|---|---|---|---|---|---|---|---|---|
|  | HMHVG | HVG | HMHVG | HVG | HMHVG | HVG | HMHVG | HVG | HMHVG | HVG |
| Virchow2 | 0.3803 | 0.1690 | **0.5561** | **0.2199** | **0.7089** | **0.3798** | **0.8717** | **0.4954** | 0.2704 | **0.1098** |
| UNI | **0.3859** | 0.1705 | 0.5221 | 0.2069 | 0.6894 | 0.3734 | 0.8714 | **0.4954** | **0.3381** | 0.1028 |
| CONCH | 0.3553 | 0.1514 | 0.4831 | 0.1808 | 0.6755 | 0.3639 | 0.8489 | 0.4724 | 0.2733 | 0.0522 |
| DINOv2-S | 0.3555 | 0.1653 | 0.4887 | 0.1722 | 0.6668 | 0.3138 | 0.8473 | 0.4717 | 0.2507 | 0.0538 |
| DINOv2-L | 0.3367 | **0.1712** | 0.4705 | 0.1822 | 0.6783 | 0.3676 | 0.8549 | 0.4798 | 0.3018 | 0.0721 |
| SAM-B | 0.2382 | 0.1111 | 0.4334 | 0.1282 | 0.6163 | 0.3211 | 0.8151 | 0.4439 | 0.2767 | 0.0947 |

### 4.8 Decoder Capacity and Objective Ablation

Table 5 compares linear, one-hidden-layer MLP, two-layer and twelve-layer attention heads trained with MSE, plus the twelvelayer Gaussian NLL reference, across five datasets and both panels. Linear and MLP outputs are unrestricted; attention heads retain softplus, so cross-family comparisons concern these practical heads rather than depth alone.

**Table 5:** Decoder and loss ablation for SimpleP2G. PCC-200 on fold 0 of five datasets, both gene panels, seed 42, with validation-selected checkpoints. Best value per dataset/panel is bold.

| Head / loss | Brain |  | PRAD |  | HER2ST |  | Lung |  | ccRCC |  | Params (M) |
|---|---|---|---|---|---|---|---|---|---|---|---|
|  | HMHVG | HVG | HMHVG | HVG | HMHVG | HVG | HMHVG | HVG | HMHVG | HVG |  |
| Linear / MSE | 0.3416 | 0.1434 | 0.5556 | 0.1888 | 0.6714 | 0.3390 | 0.8676 | 0.4732 | 0.3447 | 0.0772 | 0.512 |
| MLP / MSE | **0.3842** | 0.1426 | 0.5205 | 0.2116 | 0.6836 | 0.3518 | 0.8701 | 0.4722 | 0.3436 | 0.1025 | 1.060 |
| 2-layer / MSE | 0.3646 | 0.1518 | **0.5743** | 0.2049 | 0.6964 | 0.3763 | **0.8740** | 0.4910 | 0.3690 | **0.1115** | 16.468 |
| 12-layer / MSE | 0.3771 | 0.1670 | 0.5267 | 0.2085 | 0.7047 | **0.3878** | 0.8712 | 0.4949 | **0.3842** | 0.0866 | 45.971 |
| 12-layer / NLL | 0.3803 | **0.1690** | 0.5561 | **0.2199** | **0.7089** | 0.3798 | 0.8717 | **0.4954** | 0.2704 | 0.1098 | 45.971 |

Twelve-layer attention with MSE outperforms two-layer attention in seven of ten dataset–panel combinations, but uses 45.971 million parameters rather than 16.468 million. The MLP is strongest on Brain HMHVG (0.3842), while two-layer attention leads on PRAD and Lung HMHVG. Gaussian NLL improves over twelve-layer MSE in eight combinations, but falls from 0.3842 to 0.2704 on ccRCC HMHVG. Neither greater depth nor Gaussian NLL is uniformly preferable.

### 4.9 Gain from Spatial Context

Table 6 compares neighbors alone, the target with *K*∈ {8, 24, 48} neighbors, and the target with 24 neighbors plus a slide summary across all five datasets and both panels. The loss, decoder specification, and training protocol are fixed; changing branches also changes parameters and the initialization sequence.

**Table 6:** Conditioning-sequence ablation for SimpleP2G-cross. PCC-200 on fold 0 of five datasets, both gene panels, seed 42, with validation-selected checkpoints. Best value per dataset/panel is bold.

| **Conditioning** | **Brain** |  | **PRAD** |  | **HER2ST** |  | **Lung** |  | **ccRCC** |  |
|---|---|---|---|---|---|---|---|---|---|---|
|  | HMHVG | HVG | HMHVG | HVG | HMHVG | HVG | HMHVG | HVG | HMHVG | HVG |
| Neighbors only, *K* = 24 | 0.3159 | 0.1397 | 0.5250 | 0.2240 | 0.7203 | 0.3525 | 0.8560 | 0.4738 | 0.3903 | 0.0950 |
| Target + neighbors, *K* = 8 | 0.3779 | 0.1761 | 0.6170 | 0.2407 | 0.7263 | 0.3749 | 0.8887 | 0.5062 | 0.4080 | 0.1199 |
| Target + neighbors, *K* = 24 | 0.3804 | **0.1815** | **0.6219** | 0.2455 | **0.7352** | 0.3823 | 0.8906 | 0.5064 | 0.4157 | 0.1165 |
| Target + neighbors, *K* = 48 | 0.3797 | 0.1713 | 0.5411 | 0.2406 | 0.7279 | 0.3700 | 0.8901 | 0.5072 | **0.4171** | 0.1069 |
| Target + neighbors + slide, *K* = 24 | **0.3844** | 0.1809 | 0.5881 | **0.2460** | 0.7308 | **0.3897** | **0.8953** | **0.5109** | 0.4166 | **0.1330** |

Adding the target to the *K* = 24 neighbors-only model improves PCC-200 in all ten combinations. Adding a slide token to target+*K* 24 improves seven combinations: HER2ST HVG rises from 0.3823 to 0.3897, whereas HMHVG decreases from 0.7352 to 0.7308. PRAD HMHVG also decreases, from 0.6219 to 0.5881. Increasing neigh-borhood size is similarly nonmonotonic. Context type and tissue therefore matter beyond the amount of available image information.

### 4.10 Qualitative Analysis

We pair ResNet50 and Virchow2 on HER2ST fold 0 (SPA148) using the same SimpleP2G-cross head, training-selected gene panel and validation checkpoint rule. For target+*K* 24, replacing ResNet50 with Virchow2 increases PCC-200 by +0.0130 on HMHVG and +0.0224 on HVG. PCC improves for 137/200 and 133/199 evaluable genes, respectively. Adding a slide token changes the paired gains to +0.0187 and +0.0158. These are within-slide effects under matched training and serve a different purpose from the three-slide encoder comparison in Table 1.

Figure 3 and table 7 show the three prespecified examples: IGKC, associated with immunoglobulin expression, LYZ, associated with innate immunity, and TFF3, associated with epithelial secretion [24– 26]. The maps include all 295 measured spots with a shared color scale for each gene. In the target+*K* 24 setting, Virchow2 improves both PCC and RMSE for IGKC and LYZ. TFF3 improves in PCC, while RMSE rises from 1.5775 to 1.6090. Other transcripts can deteriorate on both metrics: LTA4H PCC decreases from 0.5930 to 0.5555 while RMSE increases from 0.4552 to 0.4724.

**Table 7:** Encoder and slide-context effects on individual genes. HER2ST SPA148, HVG, seed 42. Both contexts include the target patch. Each cell reports PCC ↑ / RMSE ↓; the best value for each metric within each gene is bold.

| **Encoder / context** | **IGKC** | **LYZ** | **TFF3** |
|---|---|---|---|
| ResNet50 / *K*24 | 0.5383 / 1.4544 | 0.8078 / 0.7931 | 0.5904 / 1.5775 |
| ResNet50 / *K*24+slide | 0.6542 / 1.3446 | 0.8043 / 0.8373 | 0.6636 / **1.3698** |
| Virchow2 / *K*24 | **0.6966** / **1.2645** | **0.8394** / **0.7474** | **0.6810** / 1.6090 |
| Virchow2 / *K*24+slide | 0.6390 / 1.3357 | 0.8347 / 0.7494 | 0.6767 / 1.4698 |

**Figure 1:**
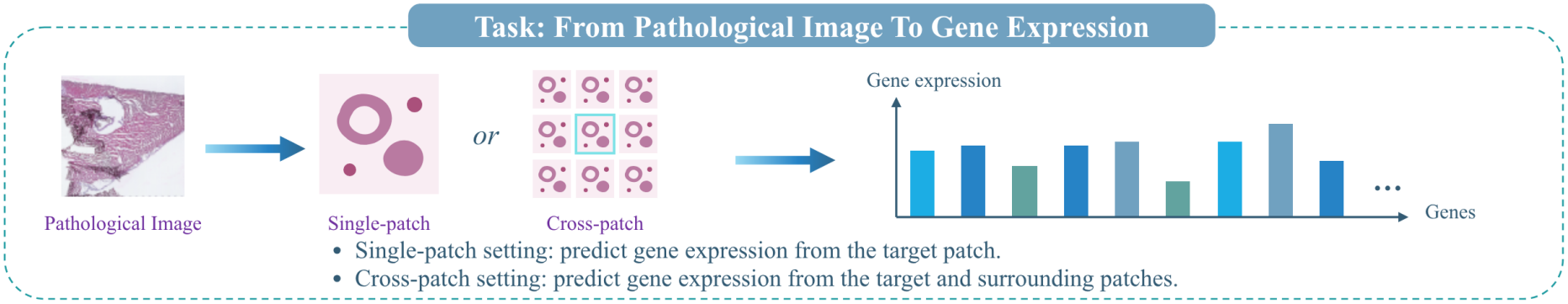
Pathology-to-gene prediction. Single-patch methods use the target patch. Cross-patch methods also use surrounding image patches.

**Figure 2:**
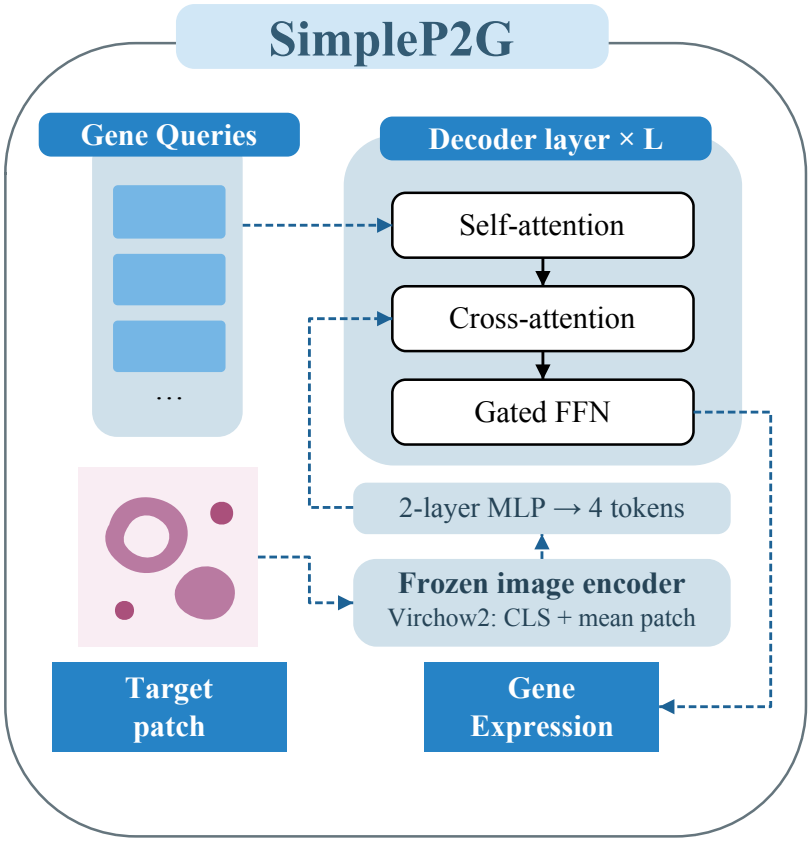
Architecture of SimpleP2G in the single-patch setting. A frozen encoder extracts patch embeddings. A small MLP converts image evidence to conditioning tokens. Learnable gene queries attend to those tokens and predict per-gene means and log-variances. SimpleP2G-cross keeps the decoder unchanged and expands only the conditioning sequence.

**Figure 3:**
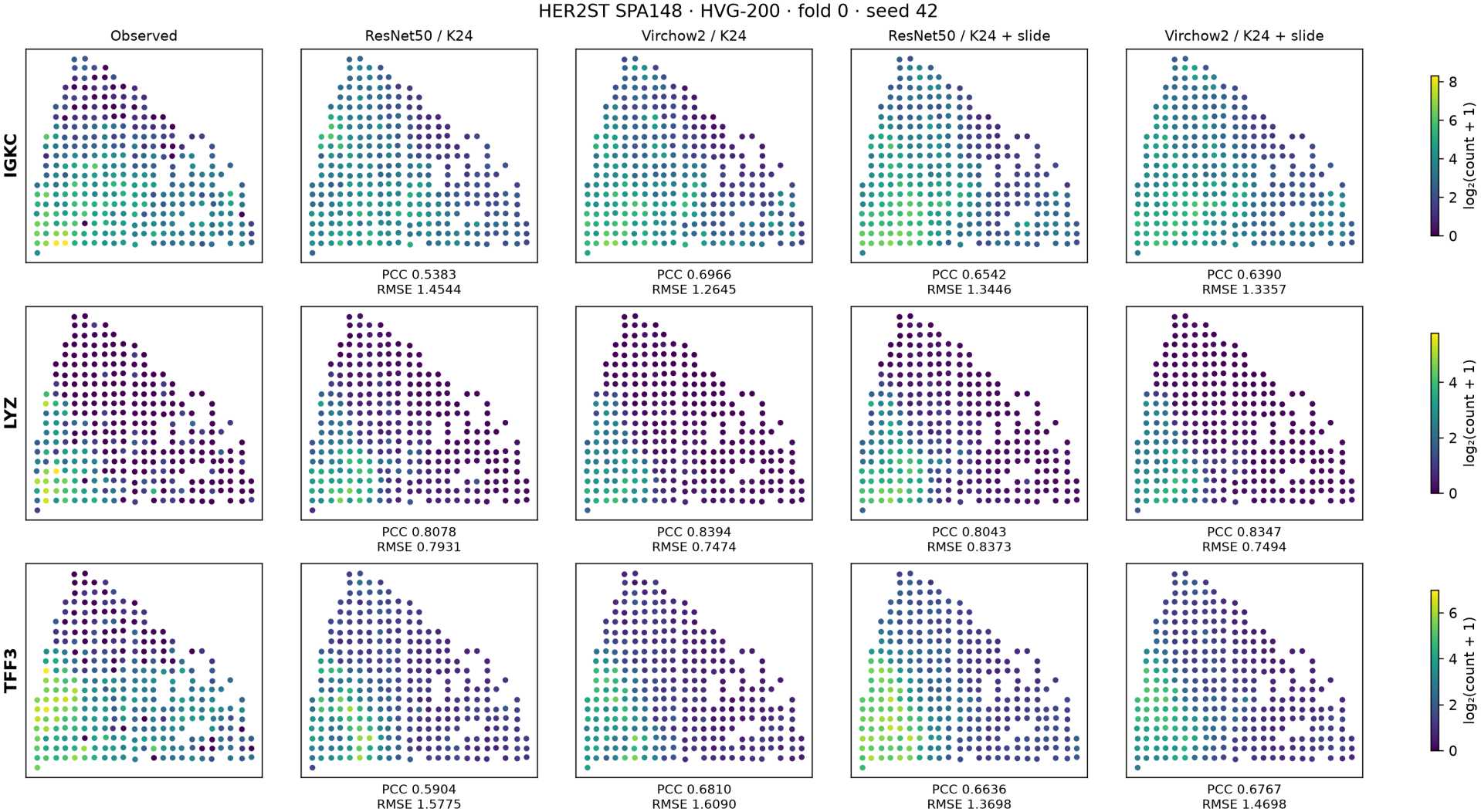
Matched encoder and context predictions on HER2ST. Observed expression and predictions for IGKC, LYZ and TFF3 on SPA148, HVG-200. ResNet50 and Virchow2 each use target+*K* 24 or target+*K* 24+slide. All 295 test spots are shown and the color scale is shared within each gene.

Adding a slide token with Virchow2 fixed decreases PCC for all three illustrated genes. For TFF3, the lower PCC (0.6810 to 0.6767) accompanies a lower RMSE (1.6090 to 1.4698). For ResNet50, slide context instead improves both metrics for IGKC and TFF3. Across the full HER2ST panels, a Virchow2 slide token improves 73/200 HMHVG genes and 108/199 HVG genes, with mean changes of −0.0044 and +0.0075. The pattern depends on the encoder, gene and metric. The exploratory Reactome test in Appendix D examines pathway associations for these changes.

## 5 Conclusion

We studied how image representation, prediction head, and spatial context affect P2G prediction. Across five datasets and both gene panels, Virchow2 improves 39 of 40 paired method–dataset–panel means. The six-encoder ablation also shows that the best representation varies with tissue and panel. Under shared Virchow2 features, SimpleP2G achieves the highest single-patch mean PCC-200 across five datasets and both gene panels. SimpleP2G-cross leads on four datasets per panel, with tissue- and panel-specific exceptions.

The decoder and context ablations show that greater depth, Gaussian NLL, and a slide token each help in several settings but are not uniformly preferable. Our central recommendation is to start with a better image representation before adding a more complex prediction head. A direct decoder over frozen pathology embeddings already provides a strong baseline against which more elaborate heads should be evaluated. These comparisons concern the specified implementations and validation-selected training protocols; the three selected slide splits do not establish patientindependent or external-cohort generalization.

We hope our reimplemented image-encoder-upgraded baselines and SimpleP2G models serve as transparent, reproducible, and strong controls for future P2G research.

## Acknowledgments

The work described in this paper was supported by the Research Grants Council of the Hong Kong Special Administrative Region, China, under Project T45-401/22-N.

## A Training and Reproducibility

The single-patch SimpleP2G settings are listed in Table 8. Both SimpleP2G variants use AdamW with *β* = (0.9, 0.999) and EMA decay 0.999. The learning rate warms up linearly and then decays on a cosine schedule to one tenth of its starting value. The experiments used PyTorch 2.10.0 and approximately 1,700 A100 GPU-hours.

**Table 8:** Dataset-specific single-patch hyperparameters. Both gene panels and all folds use the same settings.

| Dataset | LR | Weight decay | Warmup | Dropout |
|---|---|---|---|---|
| Brain | 5 × 10<sup>−5</sup> | 0.05 | 1000 | 0.05 |
| PRAD | 3 × 10<sup>−5</sup> | 0.10 | 300 | 0.10 |
| HER2ST | 10<sup>−4</sup> | 0.05 | 1000 | 0.00 |
| Lung | 5 × 10<sup>−5</sup> | 0.05 | 1000 | 0.00 |
| ccRCC | 3 × 10<sup>−5</sup> | 0.10 | 500 | 0.10 |

### Fold construction

Within each dataset, we reserve three designated test-slide IDs and choose one validation slide per fold with a fixed seed of 42. We draw uniformly from the sorted slide IDs, excluding the designated test IDs and the validation slides already assigned to earlier folds. For training, each fold uses all slides other than its own test and validation slides. The manifest records all three sets. We check gene availability against the validation and test measurement schemas and use only training expression to rank genes for panel selection.

### Precision

Most runs use FP16 mixed precision, and STRank uses FP32. Stem switches to BF16 on ccRCC fold 1, PRAD fold 1, and Brain fold 2 for HMHVG, starting at epochs 192, 130, and 158, respectively. BLEEP’s ResNet50 reference uses FP32 validation starting at epoch 99 for PRAD HVG fold 0 and epoch 87 for ccRCC HVG fold 2, and FP32 test inference for both runs. Validation scores recorded before these switches remain eligible for checkpoint selection.

### Ablation controls

The decoder/loss and context comparisons retain the split, gene panel, optimizer, learning rate, EMA, augmentation policy, and training budget of their matched reference. The twelve-layer Gaussian NLL and target+*K* 24 references reuse mainrun predictions. The conditioning variants retain the same decoder width and depth. Adding a branch introduces its own trainable projections, so the parameter count can change.

## B Baseline Implementations

The baseline training recipes use a common validation-based checkpoint rule and retain each method’s prediction objective. The details below specify the model inputs and any adaptations used in the benchmark.

### BLEEP

A contrastive loss trains the image and gene projections in a shared space of dimension 256. At inference, BLEEP retrieves the 64 most similar training spots in the projected image space and softmax-weights their expression vectors by cosine similarity. The reference trains an ImageNet-pretrained ResNet50 on 224 × 224 patches with the projection heads. In the paired run, frozen Virchow2 features supply the image projection input. Both use a learning rate and weight decay of 10^−3^.

#### Stem

The gene tokens combine a learned gene embedding with a projection of noisy counts. A 12-block diffusion transformer conditions on the target image and diffusion timestep through adaptive layer normalization, with width 384, six heads and EMA decay 0.9999. Each draw takes 1,000 diffusion steps. Validation uses one sample per spot with a fixed random stream, while test predictions average 20 samples.

#### STRank

An affine predictor is trained on raw counts with the released <monospace>stranklist_v2</monospace> ranking loss, using within-slide permutations to sample pairs. AdamW uses a fixed learning rate of 5 ×10^−5^ and weight decay 0.01. Inference receives only the target embedding. PCC compares the raw ranking scores with the common log-count targets. The scores are relative expression estimates.

#### HisToGene

An input projection and learned spatial embeddings feed a four-layer transformer over the spots of a slide. The transformer uses width 1,024 and 16 heads. The reference caches deterministic 112 × 112 RGB crops and flattens each crop into 37,632 values. The Virchow2 pair changes the input projection dimension to 2,560. Both train with MSE and a learning rate of 10^−5^.

#### STFlow

The slide-level denoiser uses spatial attention over eight neighbors and frame averaging. Runs use a zero-inflated negative binomial prior and a five-step flow sampler. We adapt the image interface to cached embeddings and use dense PyTorch batching for the 200-gene expression matrices.

#### MERGE

An eight-neighbor spatial graph is augmented with links to spatial and feature cluster centroids. Both encoder variants use the same graph construction and graph-attention predictor, with feature clusters recomputed from each encoder’s inputs. The reference uses the 256-dimensional released TCGA-pretrained ResNet18 features. A learned 2560 → 256 projection supplies the graph head for Virchow2.

#### M2TGLGO

The HEST adapter uses image and coordinate features in place of the native cell-morphology inputs. Learned gene embeddings replace the GO/Gene2Vec priors. This gives the adapter access to the shared image and coordinate inputs while changing its cell-level and gene-level information. Each training item contains up to 1,024 same-slide spots. The reference and the encoder upgrade use ResNet50 and Virchow2 features, respectively.

#### Gene-DML

The target branch receives all 261 Virchow2 tokens,the neighbor branch uses a contiguous 5 × 5 image grid, and the global branch receives the complete same-slide feature grid. Training retains target and cluster discrimination, supervised branch outputs, and their fused prediction. The cluster objective uses 18 clusters and weight 1.0. Adam uses learning rate 10^−4^ with a step decay of 0.95 every 20 epochs. The fused predictor supplies validation and test scores.

## C Image Encoder Specifications

Table 9 gives the input dimensions of the feature vectors used in the SimpleP2G encoder ablation. Virchow2’s pooled representation concatenates the 1,280-dimensional class token and the mean of its patch tokens. Gene-DML retains the token sequence for its target branch, as described in Appendix B.

**Table 9:**
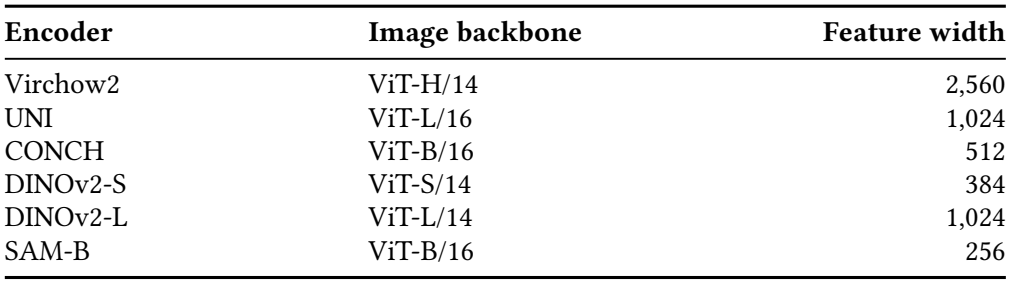
Frozen encoders in the SimpleP2G ablation. Feature width is the dimension supplied to the decoder’s image projection.

| **Encoder** | **Image backbone** | **Feature width** |
|---|---|---|
| Virchow2 | ViT-H/14 | 2,560 |
| UNI | ViT-L/16 | 1,024 |
| CONCH | ViT-B/16 | 512 |
| DINOv2-S | ViT-S/14 | 384 |
| DINOv2-L | ViT-L/14 | 1,024 |
| SAM-B | ViT-B/16 | 256 |

## D Exploratory Pathway Analysis

We test whether genes benefiting from the Virchow2 slide token on HER2ST SPA148 concentrate in particular human pathways. We match gene symbols to Reactome release 94 [20] and keep genes with variable observed expression and a pathway annotation. Of the 156 annotated HMHVG and 139 annotated HVG genes, 52 and 67, respectively, show a positive PCC change. Two-sided Fisher exact tests compare the frequency of gains inside and outside pathways with at least five background genes in each group. None of the 395 tests (283 HMHVG and 112 HVG) reaches *q <* 0.05 after joint Benjamini–Hochberg correction (minimum *q* = 0.9687).

## E Spatial Gene Expression Maps

Figure 4 shows one representative transcript per tissue across the five datasets, with one baseline and one SimpleP2G variant per input setting. Each row uses the training-selected HVG panel and validation-selected fold-0 checkpoints. Predictions preserve every test spot at its measured coordinate. All methods in a row share a color scale from zero to the observed 99th percentile of log_2_(count+1) expression. Clipping applies to the display. PCC uses the unmodified predictions.

**Figure 4:**
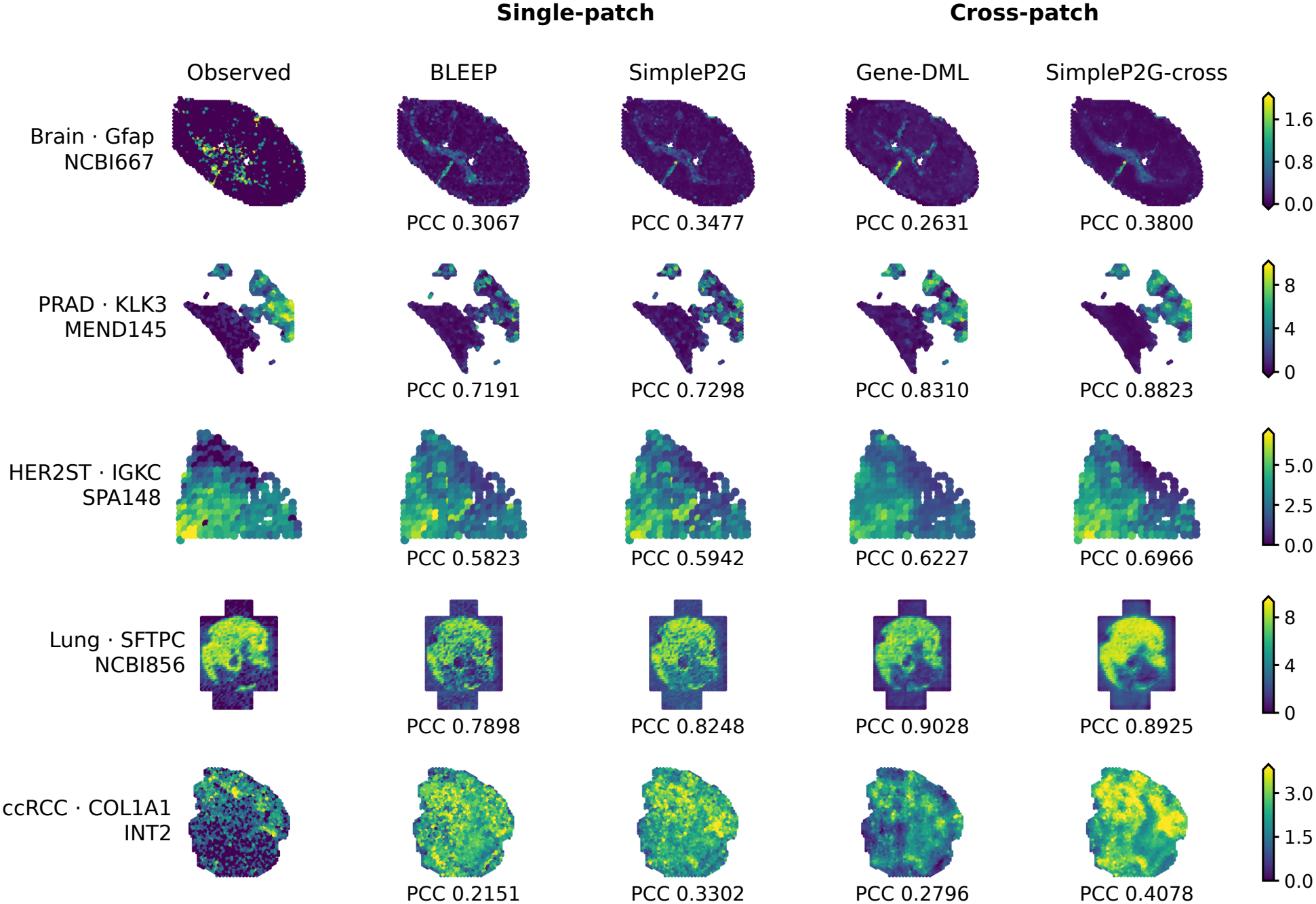
Spatial predictions across five HEST-1k datasets. Each row shows observed expression and four predictions for one gene on one held-out slide. All methods use Virchow2 features, HVG-200, fold 0, and seed 42. Per-gene PCC is printed below each predicted map.

